# Genetic and species-level biodiversity patterns are linked by demography and ecological opportunity

**DOI:** 10.1101/2020.06.03.132092

**Authors:** Chloé Schmidt, Stéphane Dray, Colin J. Garroway

## Abstract

The processes that give rise to species richness gradients are not well understood, but may be linked to resource-based limits on the number of species a region can support. Ecological limits placed on regional species richness would also limit population sizes, suggesting that these processes could also generate genetic diversity gradients. If true, we might better understand how broad-scale biodiversity patterns are formed by identifying the common causes of genetic diversity and species richness. We develop a hypothetical framework based on the consequences of regional variation in ecological limits to simultaneously explain spatial patterns of species richness and neutral genetic diversity. Repurposing raw genotypic data spanning 38 mammal species sampled across 801 sites in North America, we show that estimates of genome-wide genetic diversity and species richness share spatial structure. Notably, species richness hotspots tend to harbor lower levels of within-species genetic variation. A structural equation model encompassing eco-evolutionary processes related to resource availability, habitat heterogeneity, and human disturbance explained 78% of variation in genetic diversity and 74% of the variation in species richness. These results suggest we can infer broad-scale patterns of species and genetic diversity using two simple environmental measures of resource availability and ecological opportunity.

## Introduction

Genetic diversity and species richness are the most fundamental levels of biodiversity because they reflect within- and across-species contributions to ecosystem functioning (Oliver et al. 2015; Des Roches et al. 2021b). Genetic diversity underlies a population’s capacity to adapt in response to environmental change, and species richness enhances ecosystem resiliency to perturbation. If we are to manage the current high rates of biodiversity loss, we need to better understand how patterns of biodiversity are produced and how they interact across levels of biological organization. Patterns of species richness are well-described, but because several independent processes are capable of generating these patterns, their origins remain puzzling. We know less about multi-species patterns of genetic diversity. However, there is good reason to think that the processes forming patterns of species richness could simultaneously produce spatial patterns in neutral genetic diversity (Vellend 2005; Evanno et al. 2009). This is because spatial variation in neutral genetic diversity should reflect how local population-level demographic and evolutionary processes interact with environments to produce species richness gradients. If true, we may be able to infer processes underlying biodiversity patterns at both genetic and species levels by attempting to understand their common causes. The accumulation of open data now allows us to tackle these types of questions by repurposing and synthesizing publicly archived raw data (e.g., Leigh et al. in press; Miraldo et al. 2016; Manel et al. 2020; Schmidt et al. 2020a; Theodoridis et al. 2020; Schmidt and Garroway 2021). Here we produce a continental map of spatial variation in neutral nuclear genetic diversity for North American mammals, show that genetic diversity and species richness covary spatially and are negatively correlated, and find empirical support suggesting that measures of resource availability and heterogeneity predict both genetic diversity and species richness patterns through their effects on demography.

We developed a conceptual framework to explain how genetic diversity and species richness patterns could emerge from common causes. This framework extends predictions from well-supported hypotheses for species richness patterns to the population genetic level. Hypotheses for species richness gradients fall into three general categories related to evolutionary time, evolutionary rates, and ecological limits (Mittelbach et al. 2007; Worm and Tittensor 2018; Pontarp et al. 2019). We focus on ecological limits hypotheses—these posit that variation in resource availability limits the number of species able to coexist in a particular area (Rabosky and Hurlbert 2015). Here the speciation, extinction, and colonization dynamics of species are analogous to the birth, death, and immigration dynamics that set carrying capacities at the population level. Simulations suggest multiple hypotheses can produce species richness gradients (Etienne et al. 2019), but the preponderance of theory suggests that ecological limits produce the strongest and most stable gradients (Vellend 2005; Worm and Tittensor 2018; Etienne et al. 2019). There is also good empirical support for the likely importance of ecological limits in the formation of species richness patterns (reviewed in Rabosky and Hurlbert 2015; Brodie 2019). We thus considered ecological limits hypotheses as parsimonious starting expectations when exploring the causes of biodiversity patterns (Etienne et al. 2019).

It is relatively straightforward to extend the consequences of ecological limits on community size to the population genetic level. If environments limit the number of supportable species, they must also limit the population sizes of species, and therefore affect the strength of genetic drift. The first ecological limits hypothesis we consider is the more individuals hypothesis (Wright 1983). In terms of community composition, the more individuals hypothesis suggests that resource availability imposes an upper limit on the number of individuals, and as a consequence, the number of species an area can support (Currie 1991; Rabosky and Hurlbert 2015; Storch et al. 2018). Diversity tends to increase with the number of individuals in an assemblage both in terms of genetic diversity within populations and the number of species in a community (Kimura 1983; Hubbell 2001). Thus, the more individuals hypothesis predicts neutral genetic diversity and species richness will be positively correlated and increase with resource availability (Fig. 1).

**Fig. 1.**
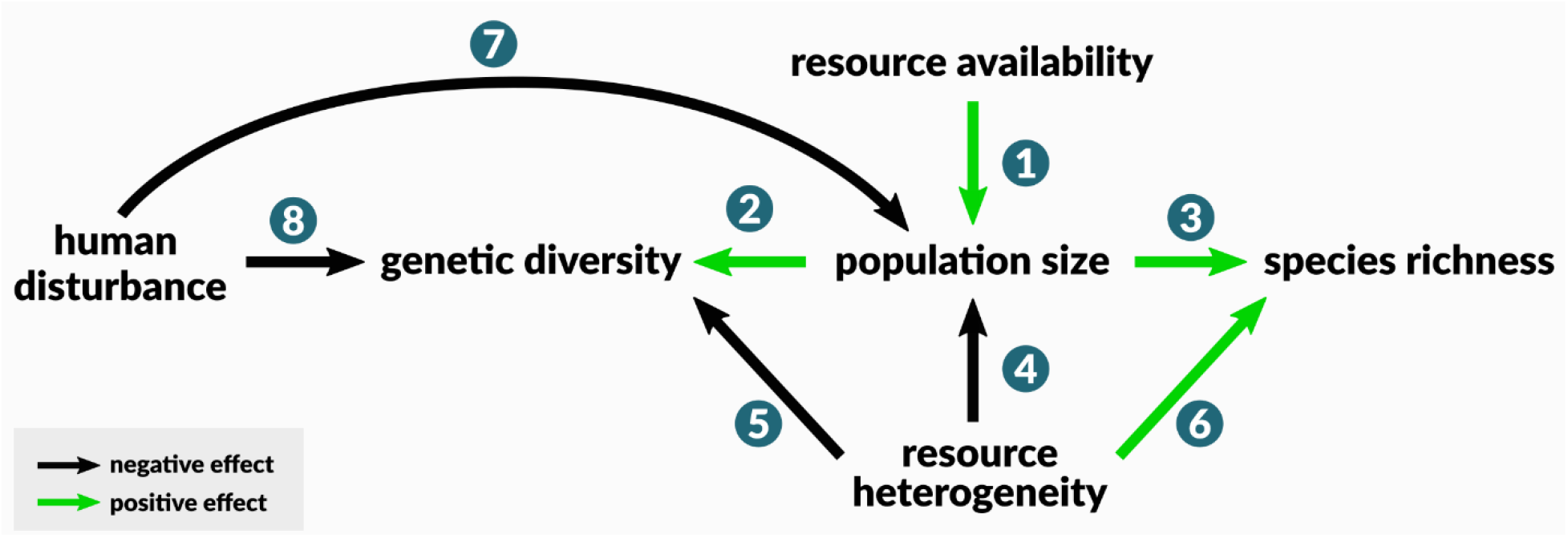
Framework integrating genetic diversity into ecological limits hypotheses. We focus on two major ecological limits pathways which stem from resource availability (the more individuals hypothesis) and resource heterogeneity. Under the more individuals hypothesis, resource availability across a species range positively affects species’ long-term effective population sizes (1). Nuclear genetic diversity increases with the effective population size (2). If population size is regulated by resource availability, it will be positively associated with community size and thus species richness (3). In heterogeneous environments, populations and species specialize to different niches but have smaller population sizes (4). Specialization reduces gene flow (5) and enhances species’ ability to coexist (6). Human land transformation reduces wildlife habitat (7) and gene flow among populations (8).

The second ecological limits hypothesis we consider pertains to environmental heterogeneity, which includes variation in resources, habitat types, and habitat complexity (Stein et al. 2014). Here we assume heterogeneity equates to niche availability. The idea of area-heterogeneity trade-offs suggests that heterogeneous areas can support richer communities of more specialized species, but these species should tend to have smaller population sizes because resources and species are divided among niches (Kadmon and Allouche 2007; Allouche et al. 2012). Local adaptation, and subsequently specialization, can also occur within and across species distributed across heterogeneous environments. As increasingly specialized populations diverge, genetic variation would be partitioned among locally adapted populations that may eventually no longer interbreed. Compared to larger populations, these smaller populations would also more rapidly lose genetic diversity due to genetic drift. If this were the case, we expect environmental heterogeneity would be positively associated with species richness and negatively associated with neutral genetic diversity (Fig. 1).

Contemporary rapid environmental change also affects biodiversity patterns, yet it is not typically modelled in a way that makes it comparable to historical processes acting over long periods. A major contemporary ecological limit on diversity is human land transformation. Human activities such as urbanization reduce the amount of habitat available to wild populations (McKinney 2006; Grimm et al. 2008) with consequences at genetic and species levels (Ceballos et al. 2015; WWF 2018; Leigh et al. 2019; Schmidt et al. 2020a). Habitat loss, fragmentation, and homogenization resulting from human land use alters resource and niche availability, thus processes associated with ecological limits should play out in populations and communities of urban wildlife. By reducing habitable area and resource heterogeneity, we predicted that the effects of urbanization for mammals should also cause species richness and genetic diversity to decrease in more heavily disturbed areas (Fig. 1).

The effects of resource availability and heterogeneity are not mutually exclusive, and in our framework they can act in concert to produce biodiversity patterns. The links among our hypotheses and their predictions are diagrammed in full in Figure 1. We jointly model both hypotheses with a method that allows us to assess their relative importance for shaping genetic diversity and species richness. Our predictions for the ways resource availability and heterogeneity interact are consistent with previous work on species richness in North American mammals (Kerr and Packer 1997), where heterogeneity becomes a more important determinant of species richness as resource availability increases. If our model successfully captures known relationships between species richness and environments, and genetic diversity behaves in the ways we predict, we will have strong empirical evidence supporting the contention that continental patterns of neutral genetic diversity and species richness are both are in part governed by ecological limits.

Our specific objectives were threefold. Because biogeographic patterns of neutral nuclear genetic diversity have not yet been mapped, we first produced a continental map of spatial patterns of genetic diversity in North American mammals. To do this we repurposed publicly archived, raw, neutral nuclear genetic data spanning 38 species and >34,000 individuals at 801 sample sites in the United States and Canada. We then tested the degree to which patterns of genetic diversity matched those of species richness. Having established shared patterns of spatial variation, we then tested our proposed conceptual model based on ecological limits hypotheses where genetic diversity and species richness are caused by common environmental factors (Fig. 1). We tested our hypothetical model using structural equation modelling (SEM), a modelling framework that fits hypothesis networks by accommodating multiple predictor and response variables. Our approach (Fig. 2) allowed us to assess the relative importance of both hypotheses and the effects of contemporary environmental change while accounting for species-level variation using hierarchical models.

**Fig. 2.**
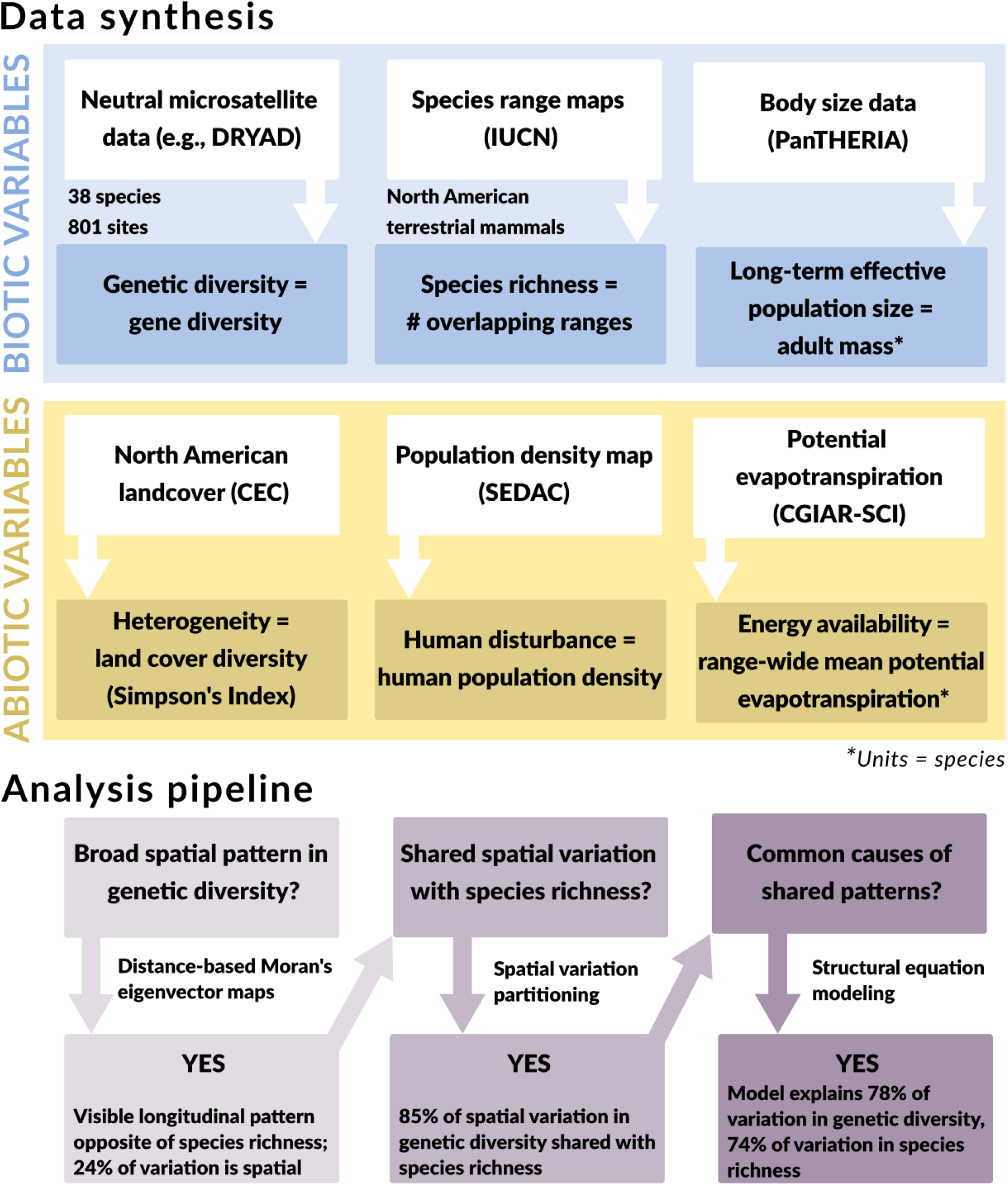
Methodological workflow detailing data sources and our series of analyses. For structural equation models, variables were either measured at each of 801 sample sites for which genetic diversity data was available, or at the species level (*n* = 38 species).

## Methods

### Biodiversity data

#### Genetic diversity

We used raw genotypic data compiled by Schmidt et al. (2020a,b). This data set is comprised of repurposed raw microsatellite data from 34,841 individuals from 38 mammalian species sampled at 801 sites in the United States and Canada. With it we could consistently calculate measures of gene diversity (Nei 1973) and population-specific F_ST_ across sites (Weir and Goudet 2017). See Table 1 for a summary of the dataset. Microsatellite markers estimate genome-wide diversity well (e.g., microsatellite and genome-wide diversity are correlated at R^2^ ∼ 0.83; Mittell et al. 2015). They are commonly used in wildlife population genetic studies because they are cost-effective and do not require a reference genome, which allowed us to maximize sample size. Detailed methods for assembling this dataset can be found in (Schmidt et al. 2020a). Briefly, we performed a systematic search for species names of native North American mammals with keywords “microsat*”, “single tandem*”, “short tandem*”, and “str” using the dataone R package, which interfaces with the DataONE platform to search online open data repositories (Jones et al. 2017). We discarded search results that did not meet our criteria for inclusion and removed results where study design may have influenced genetic diversity. For example, we excluded non-neutral data and samples taken after a recent bottleneck, translocations, managed or captive populations, or island populations. We additionally removed populations with fewer than 5 individuals sampled. Gene diversity estimates the richness and evenness of alleles in a population, and we used it here as our metric for genetic diversity because it is minimally affected by sample size (Charlesworth and Charlesworth 2010) (Fig. S1). Sample sites are treated as point locations.

**Table 1.**
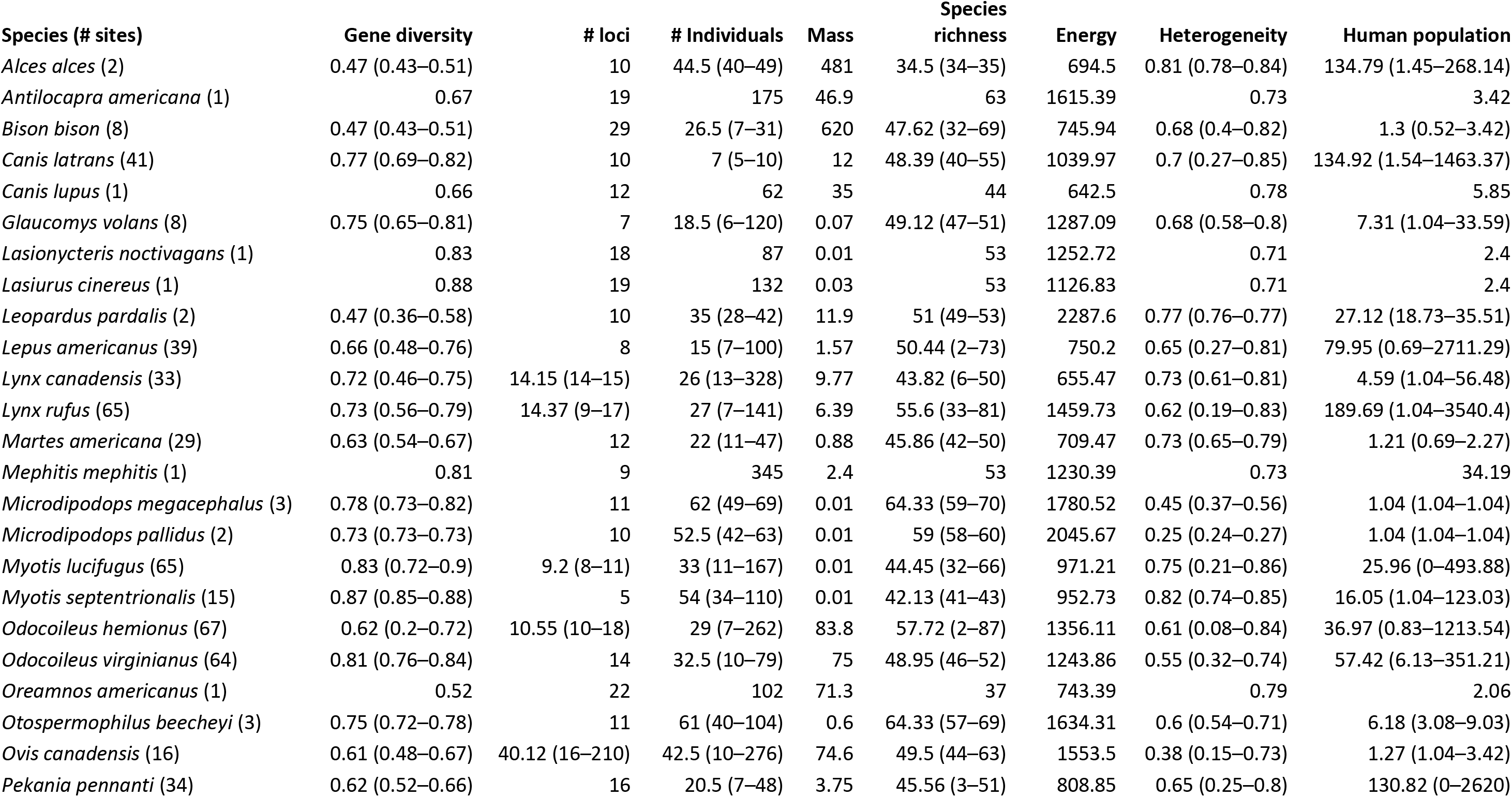
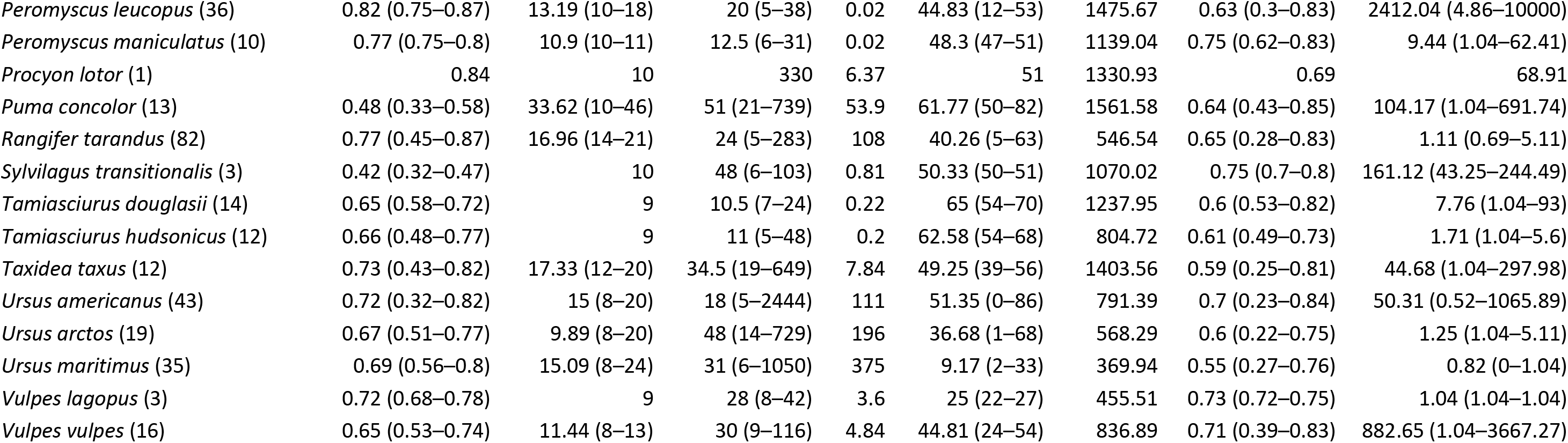
Data summary. Summary of aggregated raw genetic data: mean gene diversity, mean number of loci, median number of individuals at sites per species. Species mass (kg); species richness = mean species richness at sites; energy = mean potential evapotranspiration across species’ ranges (mm/yr), heterogeneity = mean land cover diversity (Simpson’s Index) within a 5,000 km^2^ zone around a site; human population = mean human population density across sites. Ranges of values are given in parentheses for species with multiple sample sites.

#### Species richness

We downloaded range maps for terrestrial mammals native to North America from the IUCN Red List database (IUCN 2019). We filtered these maps to retain ranges for extant, native, resident, mainland species in R version 4.0.1 (R Core Team 2020). To generate a map of species richness coincident with genetic sample sites, we estimated species richness by summing the number of ranges overlapping each site.

### Maps and spatial variation partitioning

#### Genetic diversity and species richness maps

Our first step was to map spatial patterns in genetic diversity. We accomplished this using distance-based Moran’s eigenvector maps (MEMs) in the package adespatial (Dray et al. 2017). MEMs detect spatial patterns in data using a matrix of distances between sites—a neighbor matrix—whose eigenvalues are proportional to Moran’s I index of spatial autocorrelation (Borcard and Legendre 2002; Borcard et al. 2004; Dray et al. 2006). MEMs are spatial eigenvectors that represent relationships between sites at all spatial scales detectable by the sampling scheme. Multiple MEMs can be included in linear models to identify spatial patterns in data because they are orthogonal. They are appropriate for use in genetics because Moran’s I is a direct analog of Malécot’s estimator of spatial autocorrelation of allele frequencies (Malécot 1955; Epperson 2005) which accurately summarizes neutral variation in gene flow and allele frequencies (e.g., Sokal and Oden 1978; Epperson 2005). Distance-based MEM analysis produces *n* – 1 MEMs (*n* being the number of sample sites), but only eigenvectors corresponding to positive spatial autocorrelation are used. MEMs are ordered according to spatial scale explained, with the first eigenvector explaining the broadest autocorrelation pattern. We used linear regressions and the forward selection procedure described in (Blanchet et al. 2008) to select two sets of MEMs: one describing spatial patterns in genetic diversity and the other describing species richness. Thirteen MEMs, ranging from broad to fine scales, explained important spatial variation in gene diversity. Forty-three MEMs were important predictors of species richness, and 8 of these patterns were shared by genetic diversity (significant MEMs are listed in Fig. S3).

To restrict ourselves to broad spatial patterns, we focused on genetic and species MEMs with Moran’s I values >0.25. We fit individual linear regression models for species richness and genetic diversity with corresponding broad-scale MEMs as covariates and plotted model predicted values representing spatial patterns on maps of North America (Fig. 3). These MEMs describe the broadest-scale spatial patterns at both levels of diversity. By using values of genetic diversity and species richness described by these MEMs, we can visualize pure spatial variation at the continental scale without local spatial patterns that may be due to environmental idiosyncrasies, and without considering non-spatial variation in genetic diversity. We also provided maps of raw genetic diversity and species richness values in Figure S2.

**Fig. 3.**
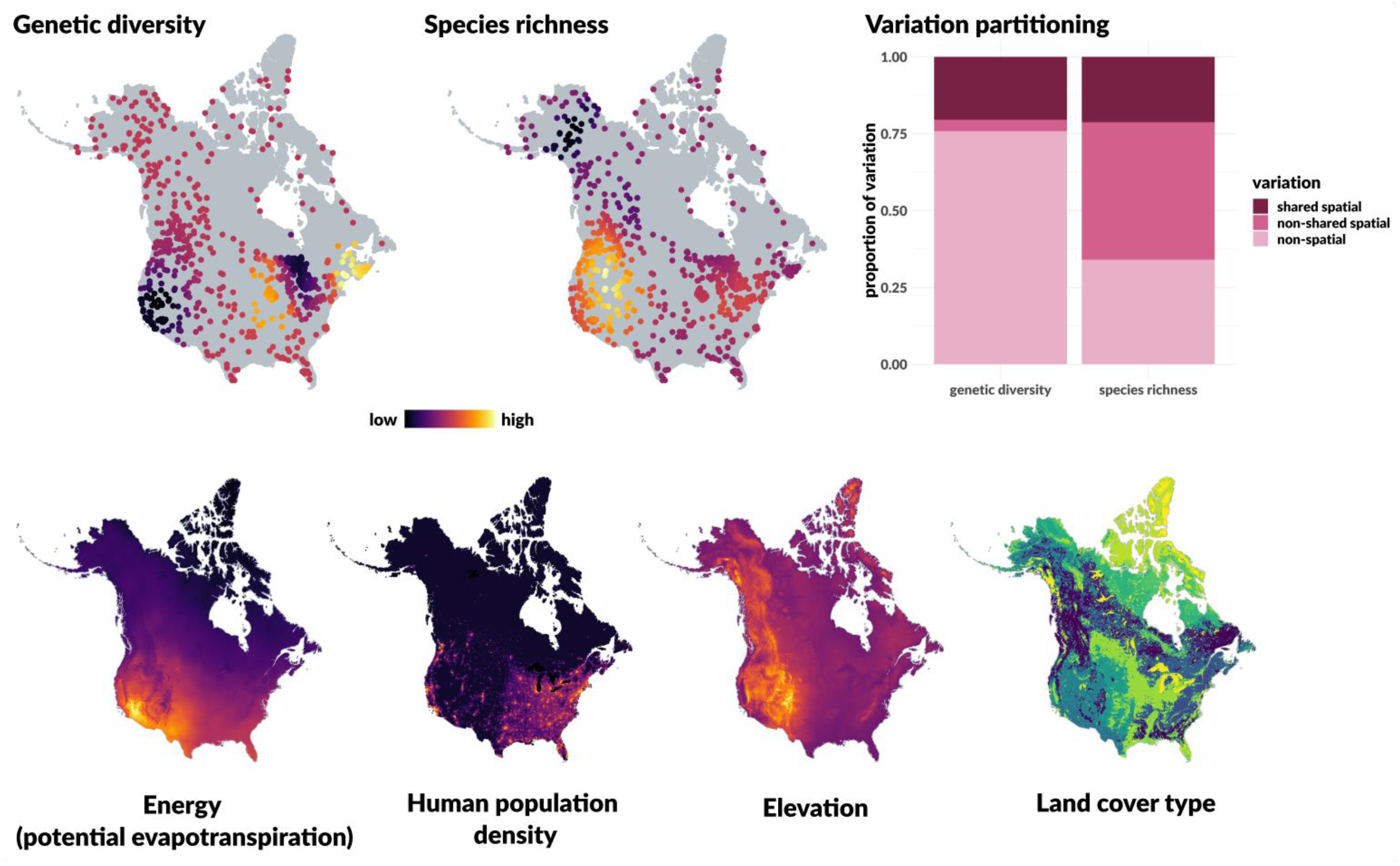
Spatial patterns of biodiversity and environmental factors. *(Top row)* Locations of 801 North American mammal populations for which raw microsatellite data was available in public repositories. Point color indicates predicted values of genetic diversity and species richness based on spatial patterns detected in the data. The variation partitioning plot shows the proportion of variation in genetic diversity and species richness which can be explained by spatial factors. Spatial variation is further broken down into shared and non-shared spatial variation. (*Bottom row*) Major environmental features of North America. Note land cover is categorical and colors represent different types. Elevation is shown for reference, but was not included in our models.

#### Variation partitioning

We next quantified the extent to which genetic diversity and species richness covary spatially. Because MEMs for species richness and genetic diversity were computed from the same set of coordinates, they were directly comparable. This allowed us to identify shared spatial MEMs. We used linear regressions and variance partitioning to determine what fraction of the total variation in species richness and genetic diversity could be attributed to: (1) non-spatial variation, (2) non-shared spatial variation, and (3) shared spatial variation. We partitioned variation as follows:

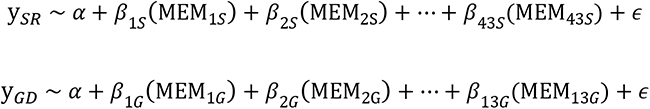

where α is the grand mean, and y_SR_ and y_GD_ are species richness and genetic diversity at sites. MEM*_S_* and MEM*_G_* refer to the set of MEMs explaining spatial variation in species richness (43 MEMs) and genetic diversity (13 MEMs), respectively, and *β*s are their slopes. The coefficients of variation (R^2^) for these models give us the proportion of variation in each response variable attributable to spatial variation. Subtracting these values from 1 gives the amount of non-spatial variation.

To determine the amount of shared variation, we used the set of 8 MEMs shared between species richness and genetic diversity (MEM_SG_) as predictors in the regressions below:

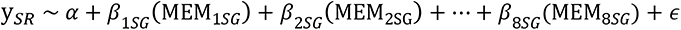

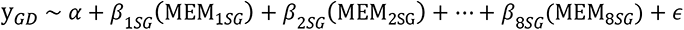

R^2^ values from these models estimate the proportion of variation in genetic diversity and species richness explained by shared spatial variation. Subtracting these values from the total spatial variation in species richness and genetic diversity gives the proportion of non-shared spatial variation.

### Structural equation modeling

#### Population size data

The more individuals hypothesis is most applicable at broad spatial scales, and when considering the total number of individuals that comprise a species (Storch et al. 2018). In place of census sizes for the species in our dataset, which are not consistently available, we craft our hypothesis around species’ long-term effective population sizes. The effective population size is a concept defined in population genetics as the number of individuals in an idealized population that experiences the same rate of genetic drift as the real population (Charlesworth and Charlesworth 2010). Populations lose genetic diversity to drift at a rate inversely proportional to the effective population size. Body size is routinely used as a proxy for long-term effective population size at the species level (Frankham 1996; Corbett-Detig et al. 2015; Mackintosh et al. 2019; Buffalo 2021). Large bodied species that tend to have long lifespans and produce few offspring generally have smaller effective population sizes than small, fecund, short-lived species (Romiguier et al. 2014; Mackintosh et al. 2019). Thus body size measured at the species level is an imperfect, but nevertheless useful substitution for census size. We recorded adult body mass (g) for each species in our genetic dataset from the PanTHERIA database (Jones et al. 2009) and log-transformed values before analysis. There were no obvious outliers in these data.

#### Environmental variables

We used potential evapotranspiration as a surrogate for total ecosystem resource availability (Currie 1991; Rabosky and Hurlbert 2015). Potential evapotranspiration is an indicator of atmospheric energy availability and is one of the strongest environmental correlates of species richness in North American mammals (Currie 1991; Kreft and Jetz 2007; Fisher et al. 2011; Jiménez-Alfaro et al. 2016). As we predict, at the species level, that resource availability across a range sets the long-term effective population size, we estimated mean range-wide potential evapotranspiration (mm/yr) using annual data from 1970-2000 available via the CGIAR Consortium for Spatial Information (Trabucco and Zomer 2019). For comparison, we also measured mean range-wide actual evapotranspiration, an alternative measure of resource availability, and present those results in the Supplementary Information.

We quantified heterogeneity and niche availability using a 250 m resolution map of land cover types in North America (CEC et al. 2010). This map includes 19 land cover categories based on satellite imagery collected in 2010 with multiple categories of forest, shrubland, grassland, polar habitat types, wetland, cropland, barren land, built up land, and open water. Because the heterogeneity hypothesis suggests species specialize on different resources within their range, we quantified heterogeneity at sites rather than at the species level. We measured heterogeneity using Simpson’s diversity index. To assess scale dependence, we calculated Simpson’s index within four buffer zones around each site: 5000, 20000, 50000, and 100000 km^2^. Lastly, we quantified human disturbance at each site using human population density (CIESIN 2016) measured within a 10 km buffer following Schmidt et al. (2020a).

#### Analysis

Structural equation modeling accommodates multiple dependent and independent variables in a model network, and directional paths connecting variables represent causal relationships. The strengths of direct paths are regression coefficients (Shipley 2016), and indirect effects can be quantified by multiplying coefficients along paths of direct effects. We constructed a graph of our conceptual model laid out in the introduction (Fig. 1), which we then translated into a network of three linear models for body size, species richness, and genetic diversity. In it, body size is predicted by resource availability, and species richness and genetic diversity are each predicted by body size, heterogeneity, and human disturbance (Fig. 4a). We fit structural equation models using piecewiseSEM in R (Lefcheck 2016; Lefcheck et al. 2019) because this package accommodates complex model structures. We used a linear mixed-effects model with a random intercept for species to account for species-level variation in genetic diversity. PiecewiseSEM fits hierarchical models using the lme4 package (Bates et al. 2015). Body size and species richness models were initially fit as linear regressions, but residuals from both models were spatially autocorrelated at broad scales. We refit these regressions using simultaneous autoregressive models in spatialreg (Bivand et al. 2013) and this successfully removed spatial autocorrelation from the residuals. All variables were scaled and centered before analysis to obtain standardized regression coefficients, allowing us to compare the strength of relationships and the relative support for hypotheses across genetic and species levels.

**Fig. 4.**
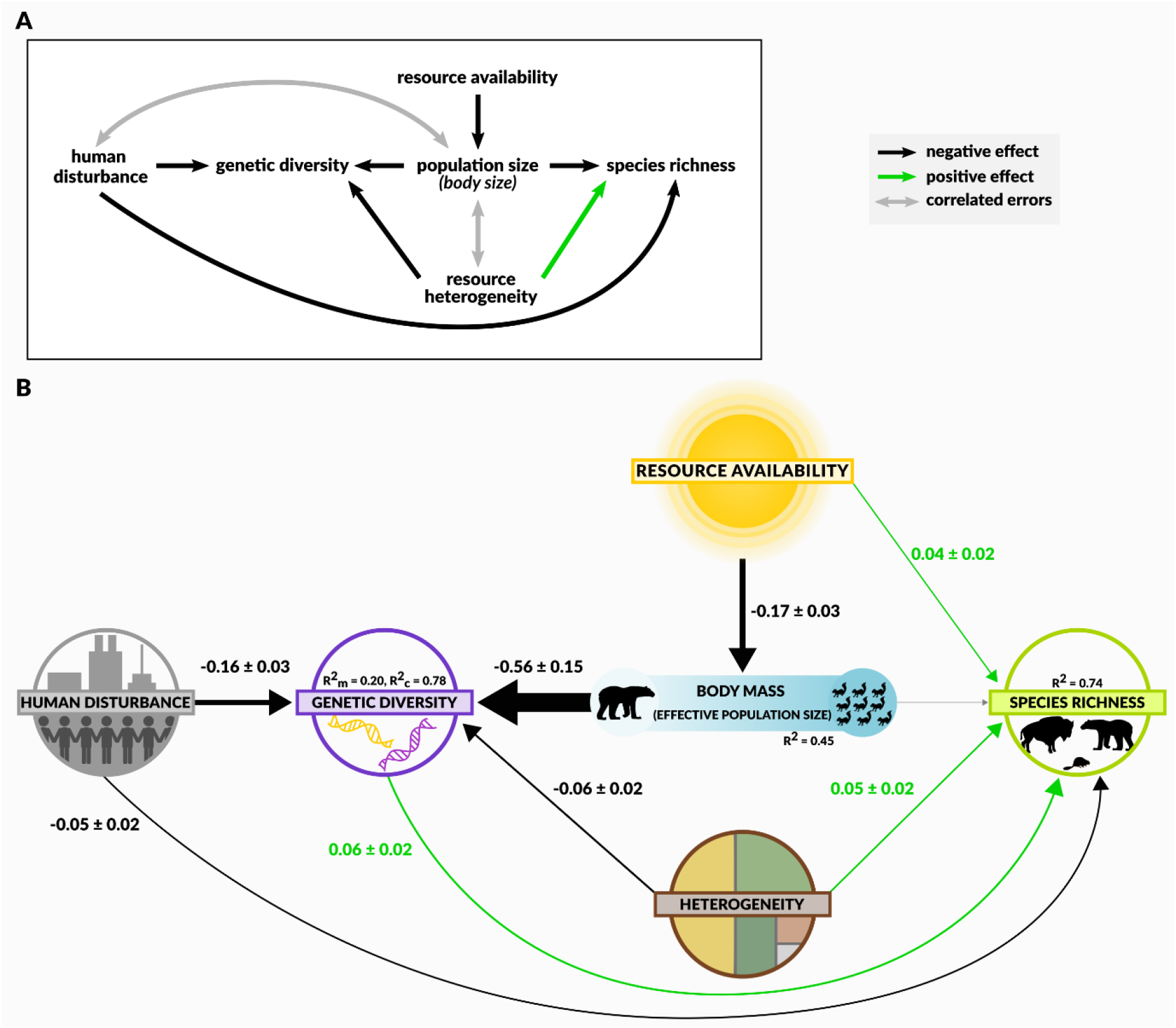
Structural equation models. (a) Our conceptual hypothesis network, modified from Figure 1 to accommodate variables measured at species and site levels. Single-headed arrows represent unidirectional relationships between variables. Grey double-headed arrows indicate variables with correlated errors that were excluded from model evaluation (see Methods). (b) Structural equation model results. Green and black lines are positive and negative relationships, respectively, and the grey line is an unsupported link. Line widths reflect partial regression coefficients, which are listed for each path with standard errors. *R^2^* values are the amount of variation explained for each response variable. Genetic diversity was fit with a random effect for species: *R^2^_m_* is the variation explained by fixed effects only, and *R^2^_c_* is the variation explained by fixed and random effects.

Our model includes variables measured at the site level (genetic diversity, species richness, heterogeneity, and human disturbance) and species level (body size, resource availability; Fig. 2). This hierarchical data structure introduces spurious correlations between variables sampled at different levels that we know are not causal. For example, regressing human disturbance at sites on species body size would estimate an artefactual experiment (the size of species researchers choose to sample near cities)—not the effects of disturbance on body size. We can account for these known non-causal relationships by allowing variables to have correlated errors (Lefcheck 2016). Correlated errors indicate that a relationship exists between variables, but allow the direction of causality to be ambiguous: both could be caused by another factor not included in the model (e.g., researcher species choice). We specified correlated errors between body size and human population density, and body size and heterogeneity.

The conceptual model is evaluated by testing whether additional links are needed between variables to make the proposed causal structure more consistent with the data. In piecewiseSEM, missing links are tested using tests of directed separation (Shipley 2016), where the null hypothesis is that two variables are independent conditional on other predictors in the model. A *p* value for the model network is obtained by testing Fisher’s C calculated from the *p* values summed across directed separation tests (Lefcheck 2016; Shipley 2016). A model-wide *p* < 0.05 means the causal structure is not a good fit to the data and additional links are needed to resolve dependencies. If *p* > 0.05, the model is considered acceptable because we fail to reject our causal structure. This means that although we start with a focus on our conceptual model, the data can suggest the addition or removal of links and our hypotheses can be updated for future testing with new data.

We assessed model fit using R^2^ values for each response variable in the model network. For genetic diversity, we used marginal R^2^ (R^2^_m_) which measures the total variation explained by fixed effects, and conditional R^2^ (R^2^) which is the variation explained by both fixed and random effects. For spatial body size and species richness regressions, we report Nagelkerke pseudo-R^2^.

### Effect of heterogeneity on population divergence

After detecting a negative effect of heterogeneity on intraspecific genetic diversity in our structural equation model, we performed a post hoc analysis to test whether environmental heterogeneity also caused greater population differentiation within species. To test for differentiation we calculated population-specific F_ST_ (Weir and Goudet 2017) as a measure of genetic divergence using the hierfstat package (Goudet and Jombart 2015). Population-specific F_ST_ can be interpreted as a relative estimate of the time since a population has diverged from a common ancestor. This metric required at least 2 sampled populations in the original studies to estimate, and due to this constraint 16 sites were excluded from this analysis. We controlled for isolation-by-distance by including MEMs significantly related to F_ST_ to account for spatial structure. We scaled and centered all variables, then used a linear mixed model controlling for species differences by including it as a random effect.

## Results

### Spatial patterns in genetic diversity and species richness

There was no obvious relationship between latitude and nuclear genetic diversity (Fig. 3). Similar to patterns of species richness, a longitudinal gradient in genetic diversity was the dominant pattern for North American mammals—however, it appears regions with high species richness have lower genetic diversity. We detected spatial patterns at genetic and species levels of diversity. Sixty-five percent of the total variation in species richness and 24% of variation in genetic diversity was spatially structured (Fig. 3). Variance partitioning suggested that 85% of the total spatial variation in genetic diversity, and 32% of spatial variation in species richness was accounted for by spatial patterns shared at both levels of diversity. This shared variation implies that, to an extent, neutral genetic diversity and species richness are simultaneously shaped by spatially structured environmental factors, and these shared processes account for most of the spatial variation in genetic diversity.

### Joint environmental causes of genetic diversity and species richness

We present results from the model using a 5000 km^2^ heterogeneity buffer in the main text. Results from wider heterogeneity buffers can be found in SI Tables S1-S4. Our conceptual model, updated according to tests of directed separation, fit the data well (SEM *p*= 0.33, Fisher’s C= 2.245; Fig. 4, Table S1). Note that for structural equation models, *p* > 0.05 indicates that we fail to reject our model. There was no residual spatial autocorrelation in body size and species richness model residuals, and genetic diversity model residuals were spatially autocorrelated at local scales (genetic diversity Moran’s I = 0.01). These Moran’s I values indicate very weak spatial structure in the data, and so we decided not to integrate it into our model. Positive spatial autocorrelation at such short distances is likely an artifact of irregular site locations and the hierarchical nature of the data. A lack of strong spatial autocorrelation in the model residuals suggests that the spatial structure of the diversity data was well captured by our model’s environmental covariates (Fig. S3). All predicted links in our conceptual model were supported except that between body size and species richness (Fig. 4). Tests of directed separation suggested additional direct links from resource availability to species richness, and genetic diversity to species richness (Fig. 4). Effects of heterogeneity on genetic diversity were not detectable at broader scales (Tables S2-S4). These relationships were consistent using actual evapotranspiration as an alternative measure of resource availability (Table S5).

Resource availability, heterogeneity, and human disturbance, acting both directly and indirectly through species population size, explained 20% of the variation in genetic diversity. The species-level variation explained by the random effect brought the total variation in genetic diversity explained by our model to 78%. The same model explained 74% of the variation in species richness. Genetic diversity was strongly negatively related to body size. The direct effects of resource heterogeneity on species richness and genetic diversity were of similar magnitude, and directions of effects were as expected if processes associated with greater resource heterogeneity reduce population sizes, lead to increased genetic drift, and facilitate local adaptation and coexistence (Fig. 4, Table S1). However, because gene diversity is not a measure of divergence, we additionally tested whether environmental heterogeneity predicted evolutionary divergence at the population level. Heterogeneity was positively related to genetic divergence, but the effect was not significant (β = 0.06 ± 0.04 SE). Finally, human disturbance negatively affected both species richness and genetic diversity, but its effects were stronger for genetic diversity (Fig. 4, Table S1).

## Discussion

We found striking continental spatial gradients in nuclear genetic diversity, and show that these patterns are negatively correlated with patterns of species richness in North America (Simpson 1964) (Fig. 3). A considerable portion of the variation in genetic diversity and species richness patterns could be explained by just three environmental factors: resource availability, resource heterogeneity, and human disturbance. This is strong empirical evidence suggesting that genetic diversity and species richness patterns emerge, in part, from the same environmental processes.

Both our maps and our structural equation model suggest that resource availability and heterogeneity interact to produce biodiversity patterns at genetic and species levels. In North America, the threshold where environmental heterogeneity presumably becomes a more important determinant of species richness than resource availability lies roughly along the US-Canada border where potential evapotranspiration reaches ∼1000 mm/yr (Kerr and Packer 1997). Near this threshold is also where we see longitudinal patterns of genetic diversity emerge. Although the negative correlation between spatial patterns of genetic diversity and species richness is most apparent in species richness hotspots (particularly in the southwest), structural equation modeling incorporating both hypotheses gives us a more nuanced view of the connections between these patterns. Indeed, effects related to resource availability and heterogeneity were of similar magnitude (Fig. 4). Population size and genetic diversity increased with resource availability, and though the link between species’ long-term effective population sizes and species richness was unsupported, species richness also increased with resource availability. Moreover, we detected a positive relationship between genetic diversity and species richness as predicted if population and community size increase with resource availability. It may be that effective population size, measured using species body size, is too coarse an indicator of census population size to detect an effect on species richness at sites—if so, site-level measures of genetic diversity could be a better indicator of local population sizes.

Our results suggest that once a minimum energy threshold is reached, populations can afford to specialize in heterogeneous environments while maintaining viable population sizes. In this way, the interplay between ecological limits and ecological opportunity simultaneously produces biogeographic patterns in genetic diversity and species richness. This interpretation of our results assumes that an environmentally set equilibrium between speciation, immigration and extinction has been reached. There is good evidence for this in North American mammals, where diversification rates have slowed as diversity increased (Alroy 2009; Brodie 2019). However, the specific ways environments shape nuclear genetic- and species-level diversity will likely differ across taxa and regions depending on whether or not they have reached equilibrium (e.g., Schmidt et al. 2021). Though we measure contemporary genetic diversity, historical variation in resource availability and heterogeneity likely contribute to the patterns we detect because they reflect whether populations have experienced large contractions in the recent past (Hewitt 2000). However, in the past when communities may not have been at equilibrium, it seems likely that other processes could have been the predominant drivers of biodiversity patterns. Indeed, hypotheses about species richness patterns have likely been a topic of debate for so long because several processes operating with different importance across the timeline of diversification are capable of producing gradients (Etienne et al. 2019). It has been suggested that time for speciation should be most detectable more immediately following broad-scale environmental change, and when all regions are colonized, habitats that provide more opportunities for speciation should over time become the most diverse (Pontarp and Wiens 2017). As diversity increases, diversification rates slow as regions approach equilibrium (Brodie 2019). It follows that the relative importance of evolutionary time and diversification rates as contributors to biodiversity patterns varies with time with patterns ultimately affected by variation in ecological limits (Rabosky and Hurlbert 2015; Pontarp and Wiens 2017; Storch et al. 2018).

Contemporary environmental change is our chance to explore pre-equilibrium processes. Cities are the newest and most rapidly expanding biome, and it is clear that they have already profoundly affected biodiversity patterns (Palumbi 2001; WWF 2018; Schmidt et al. 2020a). At this early stage of colonization it is unlikely that urban communities have reached equilibrium, suggesting processes related to evolutionary time and diversification will predominate until more niches are occupied. Indeed, there is some evidence that following an initial extinction debt after rapid urbanization, older cities support higher species richness (Aronson et al. 2014). Human disturbance had a negative effect on genetic diversity in our model, and also reduces gene flow in mammals (Schmidt et al. 2020a). This suggests that there is potential for population divergence and local adaptation if new urban niches are exploited and spatially varying selection is sufficiently strong in cities. The extent to which urban populations adapt to local environmental conditions is an ongoing and active field of study, and no consensus has been reached (Lambert et al. 2021). Equilibrium levels of genetic diversity and species richness in urban communities thus seem likely to strongly depend on resource availability and heterogeneity both within and across cities, but these aspects of urban environments are not yet well defined or understood (Norton et al. 2016; Des Roches et al. 2021a).

Notably, the negative correlation we find between spatial patterns of species richness and nuclear genetic diversity runs opposite the relatively consistent positive correlations found between species richness and mitochondrial genetic diversity gradients (Martin and McKay 2004; Adams and Hadly 2012; Miraldo et al. 2016; Manel et al. 2020; Theodoridis et al. 2020). Mitochondrial DNA has several idiosyncrasies associated with the specific biology of mitochondria that distinguish it from genetic diversity measured with neutral nuclear DNA (Schmidt and Garroway 2021). The most commonly used mitochondrial markers are the protein-coding genes *cytochrome oxidase I* and *cytochrome b*, which very likely do not evolve under neutrality (Galtier et al. 2009). Unlike neutral nuclear DNA, mitochondrial genetic diversity is not consistently related to life history, ecological traits, or census and effective population sizes (Bazin et al. 2006; Nabholz et al. 2008; James and Eyre-Walker 2020). Mitochondrial genetic diversity is thus a very different quantity than the neutral nuclear diversity estimates we use here, and its lack of relationship with population size makes it unsuited for testing hypotheses based on ecological limits. Using genetic diversity metrics estimated from neutral nuclear DNA allows us to more directly link environments to species richness through demography, population size, and by extension, species life history traits which partly set the effective population size.

Ecosystem sustainability, given environmental perturbations occurring more frequently due to human causes, depends on the resiliency of landscapes, communities, and populations (Oliver et al. 2015). Our framework and the results presented herein suggest that we can understand continental patterns of species richness and genetic diversity using two simple measures of resource availability and heterogeneity. This is potentially informative for conservation practices aiming to conserve both of these levels of biodiversity at once. Maps of neutral nuclear genetic diversity can identify regions where long-term effective population sizes may have been historically small, indicating areas where low levels of neutral genetic diversity are not necessarily of immediate conservation concern (e.g., Yates et al. 2019). However, population declines due to recent human disturbance in areas with historically low genetic diversity may warrant specific attention. Furthermore, designing protected area networks based on species richness to maintain beta diversity, or variation between sites (Bush et al. 2016; Socolar et al. 2016), will likely also capture differentiated populations with complementary genetic compositions. The connections between environments, species richness, and genetic diversity we find here suggest we should be able to make informed decisions for the joint conservation of species and genetic diversity with knowledge of few environmental parameters.

## Author contributions

C.J.G. and C.S. conceptualized the study. C.S., S.D. and C.J.G. designed the study and C.S. conducted the statistical analysis with input from S.D. and C.J.G. All authors contributed to data interpretation. C.S. wrote the first draft of the manuscript and all authors participated in editing subsequent manuscript drafts.

## Acknowledgements

We would like to thank the Population Ecology and Evolutionary Genetics group for their feedback on this manuscript. We are also grateful to the authors whose work provided the raw data for this synthesis. C.S. and C.J.G. were supported by a Natural Sciences and Engineering Research Council of Canada Discovery Grant to C.J.G. C.S. was also supported by a U. Manitoba Graduate Fellowship, and a U. Manitoba Graduate Enhancement of Tri-council funding grant to C.J.G.

## Data availability

Synthesized genetic data is available from the Dryad Data Repository (DOI: 10.5061/dryad.cz8w9gj0c). Species range boundary files and environmental data are available from open online sources (see Methods).

## Supplementary Information for

Genetic and species-level biodiversity patterns are linked by demography and ecological opportunity

**Includes:**

Figures S1-S3

Tables S1-S5

---

**Figure S1.**
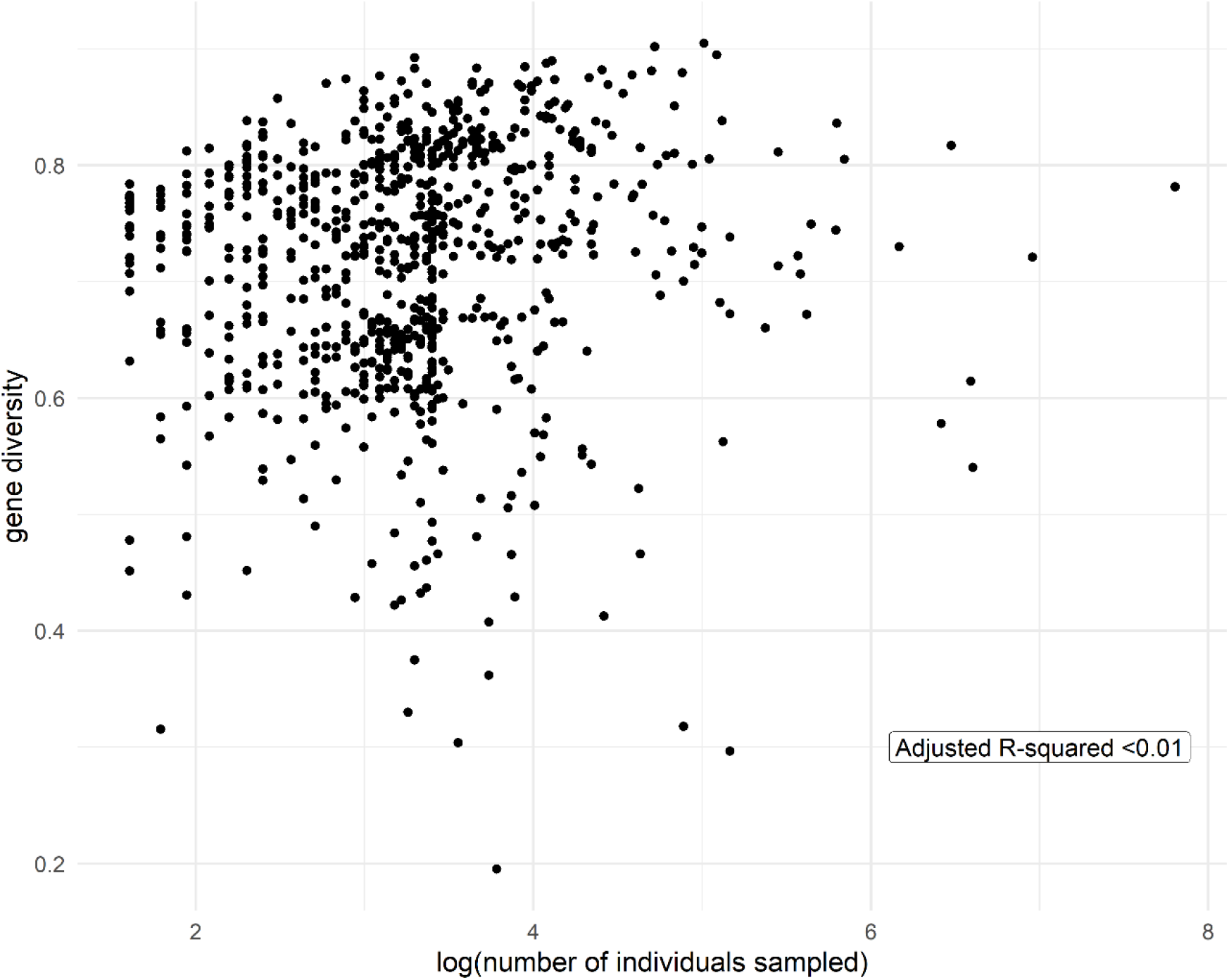
Plot of gene diversity vs. sample size. Gene diversity as a metric of genetic diversity depends on allele frequencies and is minimally affected by sample size. Larger populations have more rare alleles, which contribute little to gene diversity.

**Figure S2.**
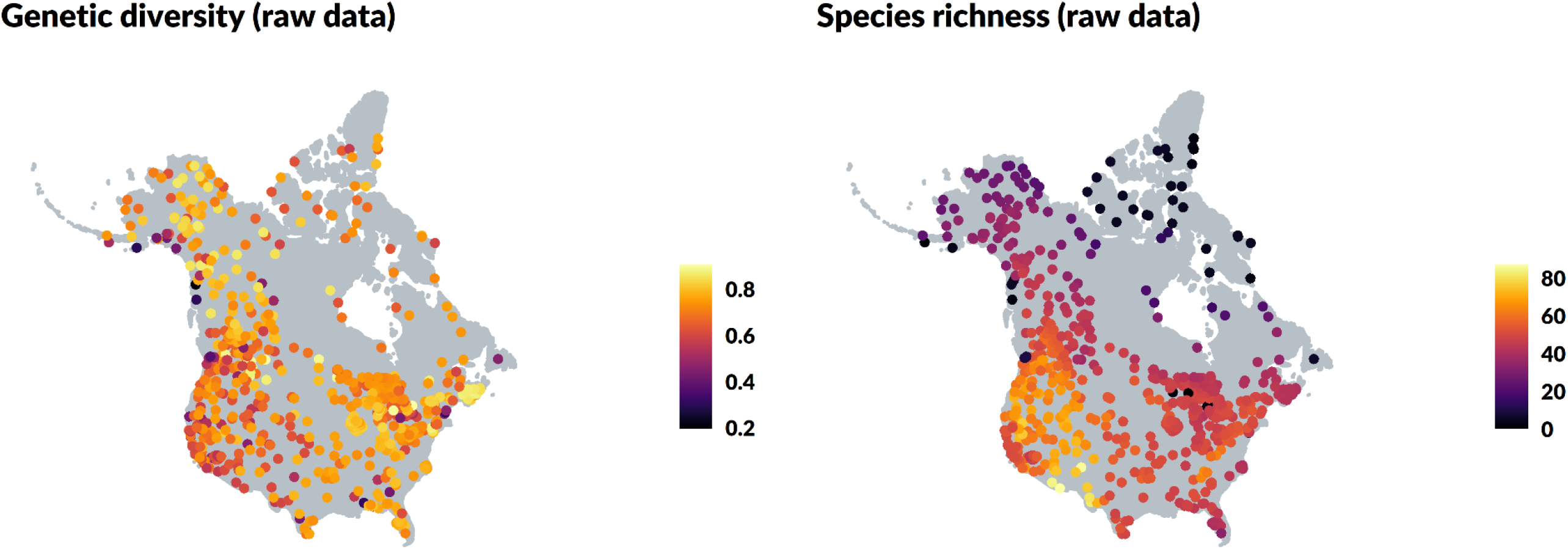
Maps of raw genetic diversity (gene diversity) and species richness data for each site. Clear gradients are visible in species richness but not genetic diversity. This is because most (65%) variation in species richness was spatial, in contrast to genetic diversity where comparatively less (24%) variation was spatially structured (Fig. 3).

**Figure S3.**
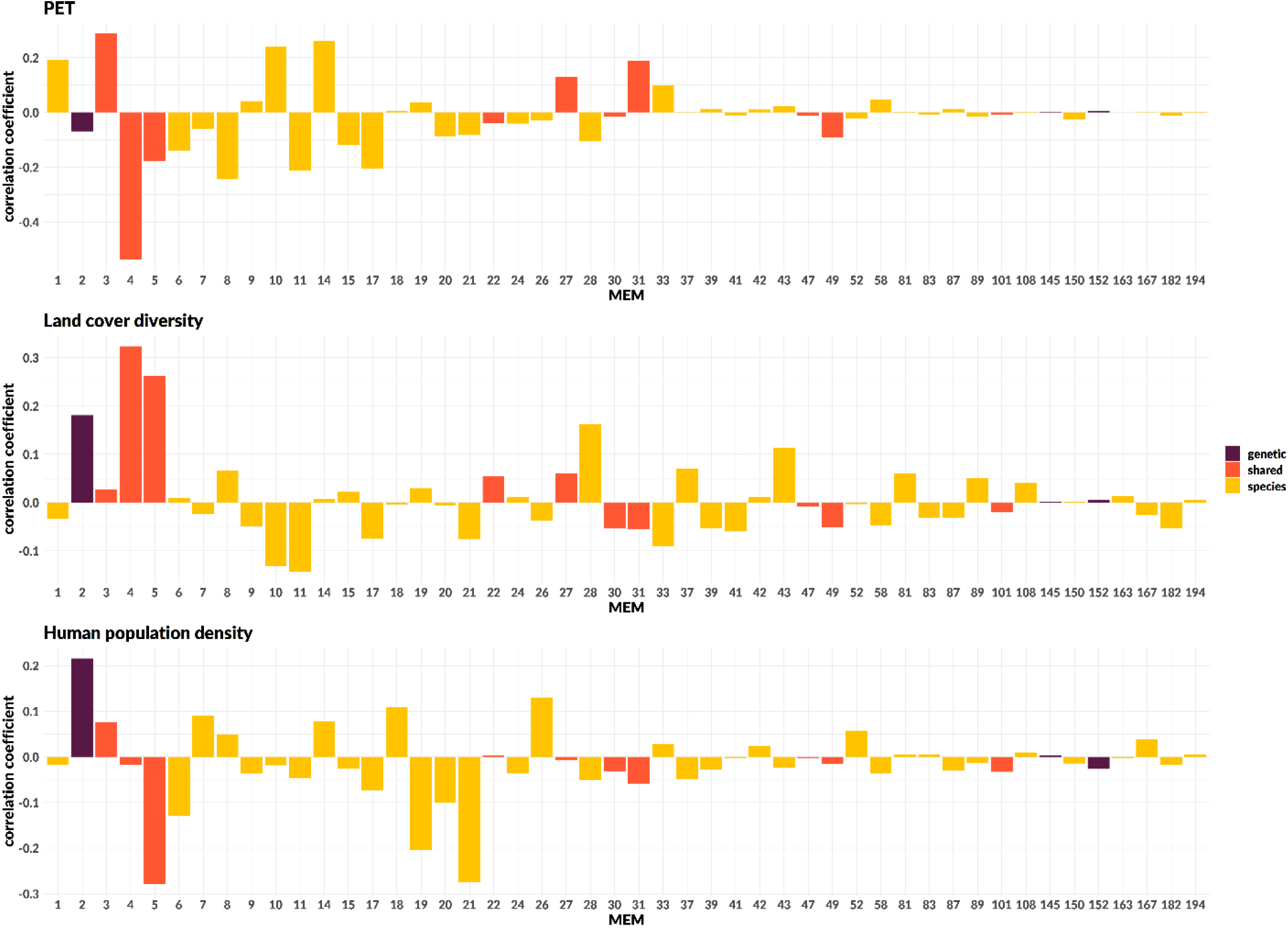
Correlation coefficients for spatial patterns (MEMs) and environmental variables measured at the site level: potential evapotranspiration (PET), land cover diversity, and human population density. MEMs describe spatial patterns in genetic diversity, species richness, or both (shared spatial patterns). MEMs are ordered from broad (MEM1) to fine scale (MEM194) patterns. Strong correlations indicate that environmental variables included in structural equation models account for broad scale spatial patterns present in genetic diversity and species richness.

**Table S1.** Path coefficients and standard errors for structural equation model presented in main text (Fisher’s C = 2.25, *p* = 0.33, 2 degrees of freedom). Heterogeneity is measured within 5000 km^2^ of a site.

| Response | Predictor | Estimate ± SE |
| --- | --- | --- |
| genetic diversity | human population density | -0.16 ± 0.03 |
| genetic diversity | body size | -0.56 ± 0.15 |
| genetic diversity | heterogeneity | -0.06 ± 0.02 |
| body size | PET | -0.17 ± 0.03 |
| species richness | body size | 0.02 ± 0.02 |
| species richness | heterogeneity | 0.05 ± 0.02 |
| species richness | human population density | -0.05 ± 0.02 |
| species richness | PET | 0.04 ± 0.02 |
| species richness | genetic diversity | 0.06 ± 0.02 |
| Correlated errors |  | Partial correlation coefficient |
| body size | human population density | 0.10* |
| body size | heterogeneity | -0.02 |

**Table S2.**
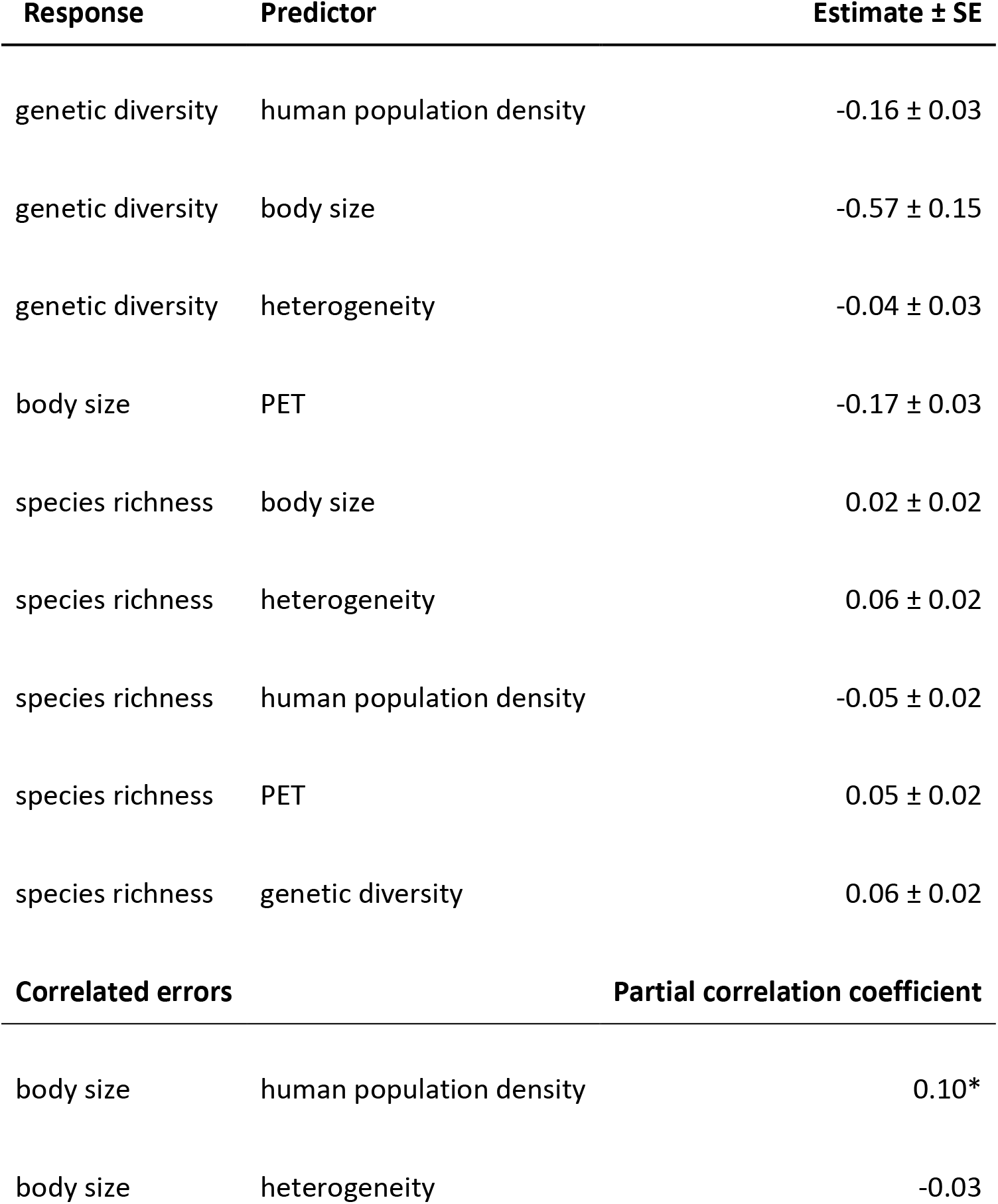
Path coefficients and standard errors for structural equation model; heterogeneity is measured within 20000 km^2^ of a site (Fisher’s C = 2.09, *p* = 0.35, 2 degrees of freedom).

**Table S3.** Path coefficients and standard errors for structural equation model; heterogeneity is measured within 50000 km^2^ of a site (Fisher’s C = 1.78, *p* = 0.41, 2 degrees of freedom).

| Response | Predictor | Estimate ± SE |
| --- | --- | --- |
| genetic diversity | human population density | -0.17 ± 0.03 |
| genetic diversity | body size | -0.57 ± 0.15 |
| genetic diversity | heterogeneity | 0.00 ± 0.03 |
| body size | PET | -0.17 ± 0.03 |
| species richness | body size | 0.02 ± 0.02 |
| species richness | heterogeneity | 0.06 ± 0.02 |
| species richness | human population density | -0.06 ± 0.02 |
| species richness | PET | 0.05 ± 0.02 |
| species richness | genetic diversity | 0.06 ± 0.02 |
| Correlated errors |  | Partial correlation coefficient |
| body size | human population density | 0.10* |
| body size | heterogeneity | -0.03 |

**Table S4.** Path coefficients and standard errors for structural equation model; heterogeneity is measured within 100000 km^2^ of a site (Fisher’s C = 1.54, *p* = 0.46, 2 degrees of freedom).

| Response | Predictor | Estimate ± SE |
| --- | --- | --- |
| genetic diversity | human population density | -0.18 ± 0.03 |
| genetic diversity | body size | -0.56 ± 0.15 |
| genetic diversity | heterogeneity | 0.04 ± 0.03 |
| body size | PET | -0.17 ± 0.03 |
| species richness | body size | 0.02 ± 0.02 |
| species richness | heterogeneity | 0.05 ± 0.02 |
| species richness | human population density | -0.05 ± 0.02 |
| species richness | PET | 0.04 ± 0.02 |
| species richness | genetic diversity | 0.06 ± 0.02 |
| Correlated errors |  | Partial correlation coefficient |
| body size | human population density | 0.10* |
| body size | heterogeneity | -0.03 |

**Table S5.**
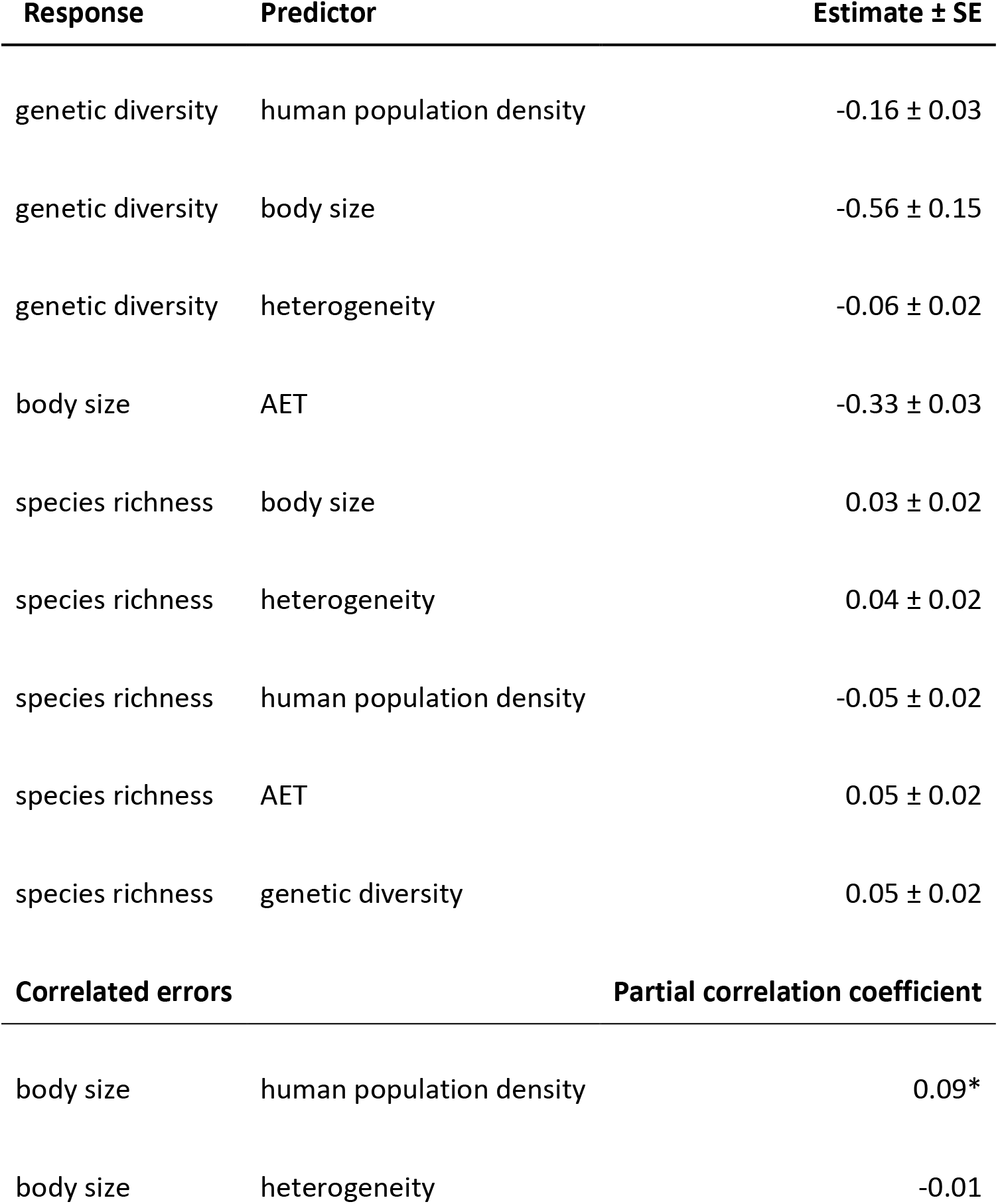
Path coefficients and standard errors for structural equation model using actual evapotranspiration (AET) as a measure of resource availability. Heterogeneity is measured within 5000 km^2^ of a site (Fisher’s C = 0.53, *p* = 0.77, 2 degrees of freedom). Genetic diversity R^2^_m_ = 0.20; R^2^_c_ = 0.78; body size R^2^ = 0.48; species richness R^2^ = 0.74.

## Notes

https://datadryad.org/stash/dataset/doi:10.5061/dryad.cz8w9gj0c

### Competing Interest Statement

The authors have declared no competing interest.

